# Evidence from three taxonomically distinct species for a non-AhR mechanism of developmental neurotoxicity of an environmentally derived mixture of polycyclic aromatic hydrocarbons

**DOI:** 10.64898/2026.08.12.743995

**Authors:** Samantha E. Phelps, Melissa Chernick, Javier Huayta, Aliyah Webster, Abigail S. Joyce, Kerry M. Ettinger, Caroline Beggs, Safiatou Zibo, Lee Ferguson, Richard T. Di Giulio, Joel N. Meyer, Nishad Jayasundara

## Abstract

Typical environmental exposures to the toxic class of chemicals known as polycyclic aromatic hydrocarbons (PAHs) involve complex mixtures; however, relatively few mechanistic toxicity studies have evaluated them as environmental mixtures, instead focusing on individual compounds or simple mixtures. In this study, we first derived Republic Sediment Extract (REPSE), a complex PAH mixture extracted from sediment at the Republic Creosoting site of the Elizabeth River in Norfolk, Virginia. After characterizing the PAH contents of REPSE, we evaluated its mechanisms of developmental neurotoxicity in three evolutionarily distinct taxa, leveraging the unique strengths of Atlantic killifish, zebrafish, and *Caenorhabditis elegans* as model species, with a focus on the Aryl hydrocarbon Receptor (AhR) pathway. Embryonic REPSE exposure caused induction of CYP1A in both fish species at sub-teratogenic concentrations, consistent with activation of the canonical AhR pathway. These sub-teratogenic exposures nevertheless induced neurotoxicity across both fish species, altering neurobehavioral phenotypes in fish, and induced dopaminergic neuronal damage in worms, again at non-teratogenic concentrations. To determine whether these effects were linked to canonical AhR response pathways, we examined killifish offspring from the pollution-adapted Republic Creosoting population, which exhibited characteristic recalcitrance to CYP1A induction, but remained susceptible to the neurobehavioral effects of REPSE. The induction of neuronal damage in worms provides orthogonal evidence for a non-AhR mechanism, because *C. elegans* AhR is not transcriptionally activated by PAHs as in vertebrates. Further probing of potential mechanisms underlying REPSE-induced neurotoxicity in worms revealed altered neuronal redox status (roGFP) and energy availability (ATP:ADP ratio). Collectively, our multispecies approach reveals conserved mechanisms of PAH mixture neurotoxicity, including effects that extend beyond canonical AhR signaling.

## 1. Introduction

### 1.1. Mechanisms of Complex PAH Mixture Toxicity

Environmental exposures are rarely limited to single contaminants. Because organisms are continuously exposed to complex chemical mixtures, understanding mixture effects is crucial to predicting outcomes. Creosote is a chemically complex mixture used by wood treatment facilities and is an environmental contaminant in the Elizabeth River in Norfolk, Virginia, a region with a long history of creosote use lasting through the late 20^th^ century (Di Giulio and Clark 2015). Up to 85% of creosote composition consists of combustion byproducts known as polycyclic aromatic hydrocarbons (PAHs) (Mueller et al. 1989). PAHs are associated with a wide variety of adverse health outcomes, including carcinogenesis, reproductive toxicity, birth defects, immunotoxicity, and neurodevelopmental disorders (Moorthy et al. 2015; Olasehinde and Olaniran 2022; Humphreys and Valdés Hernández 2023). Some but not all aspects of chronic PAH toxicity are mediated by the Aryl hydrocarbon Receptor (AhR), a ligand-activated transcription factor (Incardona et al. 2011; Van Tiem and Di Giulio 2011; Clark et al. 2010). AhR pathway activation induces xenobiotic metabolism pathways, including cytochrome P450 1A (CYP1A) enzymes, which generate mutagenic metabolites (Rotchell et al. 2008). The AhR-mediated mechanism, however, does not fully capture PAH toxicity (Billiard et al. 2008).

Despite evidence that the mechanisms of toxicity fundamentally differ between PAH mixtures and individual PAHs (Slotkin et al. 2017), mechanistic studies of environmentally relevant PAH mixtures remain limited. Sediment extracts derived from polluted sites offer a way to investigate this using a real-world mixture. In this study, we investigated sediment extracts derived from PAH-contaminated Superfund sites in the Elizabeth River in Norfolk, Virginia. We generated and characterized the contents of a novel sediment extract from the unremediated Republic Creosoting site, which we will refer to as Republic Sediment Extract (REPSE). We then used a multispecies approach to evaluate the developmental neurotoxicity of REPSE in three evolutionarily distinct model organisms: Atlantic killifish (*Fundulus heteroclitus*), zebrafish (*Danio rerio*), and nematode worms (*Caenorhabditis elegans)*. We also utilized Elizabeth River Sediment Extract (ERSE), porewater that was created prior to the remediation of the Atlantic Wood Industries Superfund Site (Fang et al. 2014), in nematode worms.

### 1.2. Atlantic killifish (*Fundulus heteroclitus*) as an ecologically relevant model organism

Atlantic killifish are uniquely relevant to the study of PAH mixtures. This estuarine species has rapidly and repeatedly evolved resistance to chemical contamination in their habitat (Reid et al. 2016). Their small home range, non-dispersive spawning, and short generation time contributes to this rapidity (Burnett et al. 2007; Lotrich 1975), allowing subpopulations to adapt to site-specific contaminants. For this reason, they have been extensively used to study environmentally relevant mixtures in the context of pollution adaptation.

The southern branch of the Elizabeth River, VA harbors subpopulations of killifish that have evolved resistance to PAH toxicity (Di Giulio and Clark 2015). Some aspects of this resistance are driven by suppression of the AhR signaling pathway and its downstream targets (Wills et al. 2010; Clark et al. 2010). Other characteristics of PAH resistance include altered redox metabolism (Meyer et al. 2003; Bacanskas et al. 2004) and mitochondrial bioenergetics (Du et al. 2016; Jasperse et al. 2023). Here, we compared the effects of REPSE in PAH-sensitive killifish to PAH-adapted Elizabeth River killifish from the unremediated Republic Creosoting site. Comparing two populations with divergent AhR pathway inducibilities allowed us to gain mechanistic information regarding the role of AhR in REPSE toxicity.

### 1.3. Zebrafish (*Danio rerio*) as a model organism for developmental toxicity

Zebrafish are an important *in vivo* vertebrate model for mechanistic toxicology studies of environmental mixtures. Many AhR agonists, including PAHs, are known to cause developmental neurotoxicity in zebrafish, as reflected by changes in neurobehavioral phenotypes such as locomotor activity and learning (Glazer et al. 2016; Stickler et al. 2024; Hawkey et al. 2022). While AhR signaling may play a role in the neurodevelopmental toxicity of individual PAHs and manually curated mixtures (Knecht et al. 2017; Geier et al. 2018), whether neurotoxicity of environmentally derived PAH mixtures is AhR-dependent has not been definitively shown. We took advantage of a transgenic zebrafish line that allowed us to quantify *cyp1a* expression as a biomarker for AhR activation following exposure to sediment extracts (Kim et al. 2013). This, along with locomotor activity assessments to evaluate neurotoxicity (Wilson et al. 2023), allowed us to evaluate the relationship between AhR activation and neurodevelopmental toxicity.

### 1.4. *C. elegans* as a mechanistic model organism

Nematode worms (*C. elegans*) are a well-characterized model organism and have been used to determine the effects of exposure to PAHs (Harris et al. 2020; Polli et al. 2020), including effects on neurodevelopment and dopaminergic neurons (Huayta et al. 2025). Dopaminergic neurons are particularly relevant because environmental toxicant exposure has been linked to neurotoxicity and neurodegenerative diseases such as Parkinson’s disease, which is characterized by degeneration of dopaminergic neurons in the substantia nigra (Dorsey and Bloem 2024). Therefore, we used *C. elegans* to quantify neurotoxic effects after exposure to sediment extracts (REPSE and ERSE), focusing on dopaminergic neuronal damage and its associated molecular phenotypes.

Mitochondria are particularly sensitive to the effects of exposure to PAHs, and mitochondrial dysfunction is a common driver of neurotoxicity due to neurons’ high energy demand (Meyer et al. 2018; Moreira et al. 2010). The transparent body of *C. elegans* enables the use of fluorescent reporters *in vivo* to quantify mitochondrial homeostasis and link it to the loss of neuronal integrity. Fluorophores such as roGFP and Perceval allow for the quantification of redox state and ATP levels, respectively, with high-spatial resolution (Morton et al. 2023). However, PAHs do not bind to and activate AhR in *C. elegans* as they do in vertebrates (Powell-Coffman et al. 1998; Butler et al. 2001) and worms do not endogenously possess the family 1 P450s that mediate much of PAH metabolism in vertebrates (Harlow et al. 2018; Leung et al. 2010). Because these pathways are absent, C. elegans provide a system for identifying toxic effects that occur independently of canonical vertebrate AhR signaling and CYP1-mediated PAH metabolism.

### 1.5. Study Objectives

The objective of this study was to evaluate the role of AhR as a mechanism of developmental neurotoxicity resulting from exposure to an environmentally relevant PAH mixture. We harnessed the strengths of three taxonomically distinct species to characterize the developmental neurotoxicity of a newly generated sediment extract, REPSE, and investigate the mechanisms underlying its effects. By using complementary model organisms, we sought to evaluate the relative contributions of AhR-dependent and AhR-independent pathways to REPSE neurotoxicity. A comparison of PAH-sensitive to PAH-adapted Atlantic killifish provided ecologically relevant information in a subpopulation of fish that had developed resistance to this PAH mixture *in situ*. Transgenic zebrafish and *C. elegans* were used to further our mechanistic understanding of AhR and how it relates to neurotoxicity. Zebrafish have canonical PAH-responsive AhR signaling that is both well-defined and measurable, whereas *C. elegans* lack this pathway, allowing us to determine AhR-independent mechanisms. In addition, because neurotoxicological effects of the previously characterized Elizabeth River sediment extract (ERSE) have not been examined in *C. elegans*, we included ERSE in the worm studies to compare mechanistic responses between the two environmentally derived PAH mixtures. We hypothesized that REPSE would cause developmental neurotoxicity across taxa, and that REPSE neurotoxicity would be associated with CYP1A activation in those species with PAH-inducible AhR pathways.

## 2. Methods

### 2.1. Creation and Characterization of Republic Sediment Extract (REPSE)

#### 2.1.1. Sediment Collection and Preparation of Sediment Extracts

River sediment was collected at low tide adjacent to the former Republic Creosoting facility in the Elizabeth River, VA, USA (36°47’35.4”N, 76°17’38.6”W) in July 2023. The top ≈30 cm of river sediment was taken from a point determined by preliminary sediment sampling, previous reports, and proximity to perceived killifish egg laying locations (Clark et al. 2013; Volkoff et al. 2019). Two 5-gallon plastic buckets were filled to the top with sediment and sealed with a lid to prevent loss of volatile compounds during transport. Buckets were transported to the laboratory at Duke University (Durham, NC, USA), and kept at 4 °C for no longer than one week prior to processing to avoid excessive PAH-to-plastic adsorption.

Sediment extract was prepared according to previous methods (Clark et al. 2013; Fang et al. 2014; Volkoff et al. 2019). Throughout, sediment and extract were protected from light whenever possible. In batches, wet sediment was combined in an approximately 1:1 ratio with deionized (DI) water, mixed vigorously, then allowed to settle. The resulting water layer was decanted into a clean bucket, siphoned into 50 mL centrifuge tubes (VWR International, Radnor, PA, USA), and centrifuged at 4300 *g* for 5 min (Sorvall RT7, ThermoScientific, Waltham, MA, USA) to remove fine particulates. Supernatant from all tubes was then decanted into a single large container, where it was continuously agitated while being siphoned into ≈40 mL aliquots. Aliquots were immediately frozen and stored at −80 °C away from light until use. Hereafter, this is referred to as Republic Sediment Extract (REPSE). The preparation and analysis of the Elizabeth River Sediment Extract (ERSE) from Atlantic Wood sediment (36°48′27.2′′N, 76°17′38.1′′W) was published previously by Fang et al. (2014). ERSE generated for that study was stored at −80 °C away from light until use.

#### 2.1.2 Determination of PAHs in REPSE

PAH porewater concentrations were measured using solid-phase extraction (SPE) and analyzed by gas chromatography combined with electron impact mass spectrometry (GC-EIMS). Two aliquots of REPSE were thawed in cold water and vortexed (n=2/REPSE aliquot). Sample blanks (DI water; n=2) underwent extraction at the same time. SPE tubes (Discover DSC-18, Supleco Inc., Bellefonte, PA, USA) with PTFE frits (20 µm porosity, Supleco Inc.) were placed in an SPE manifold and conditioned with hexane followed by acetone. Samples (10 mL) were placed in each tube and spiked with a mass-labeled surrogate standard mix containing six deuterated PAH standards: d_10_-2-methylnaphthalene, d_10_-Pyrene, D_12_-chrysene, d_12_-Perylene, D_12_-indeno(1,2,3-c,d)pyrene, and d_10_-Phenanthrene. The recoveries of these standards were only used for quality assurance/control and were not used to correct PAH values.

Samples were eluted from SPE using 1:1 acetone:hexane (6 mL) and concentrated using evaporation under nitrogen gas (N_2_; Turbo Vap, Caliper LifeSciences, Hopkinton, MA, USA) to approximately 0.5 mL. Then they were transferred to labeled amber glass autosampler vials and spiked with a deuterated internal standard mix (100uL, d_8_-Naphthalene, d_10_-Anthracene, d_10_-Fluoranthene, D_12_-benz(a)anthracene, d_12_-Benzo(a)pyrene, and D_14_-Dibenzo(a,i)pyrene). Samples were analyzed by gas chromatography mass spectrometry (Agilent GC 6890N, MS 5975, Newark, DE, USA). Data was processed using ChemStation (Agilent, Newark, DE, USA) and quantified using a 7-point internal standard calibration curve. Method details have been previously reported (Fang et al. 2014). Recoveries of PAH standards ranged from 57% to 84% except for d_12_-Perylene, for which recoveries were 32%.

### 2.2. Killifish (*Fundulus heteroclitus*) Methods

#### 2.2.1 Killifish: Collection and Breeding

All capture, care, reproductive, and experimental techniques were approved by the Duke University Institutional Animal Care and Use Committee (A069-22-04 and A069-22-04-25). Adult killifish were collected from the PAH-adapted Republic (REP) population (36°47’34.8”N 76°17’38.5”W) and a reference site at King’s Creek (KC), a relatively uncontaminated tributary of the Severn River, VA, USA (37°18′16.2″N, 76°24′58.9″W). Wire mesh minnow traps were used to capture adult fish that were sorted by sex and size on site and transported to Duke University, Durham, NC, USA. Fish were maintained in flow-through systems containing 10-15 ppt artificial saltwater (ASW; Instant Ocean, Pentair Aquatic Ecosystems, Apopka, FL, USA) and fed pelleted feed twice daily (Aquamax® Fingerling Starter 300, Lakeway Tilapia). Systems were maintained at 25-28 °C with a 14:10 light:dark cycle.

The experiments in this study were performed using offspring bred from adult killifish collected in this manner from 2022-2025. Fish were bred two weeks apart on the new or full moon. Each breeding event was considered an independent experiment. During these breeding events, gametes were obtained by manual spawning. Eggs were fertilized *in vitro* by mixing them with sperm in a beaker containing 300 mL 25 ppt ASW for 1 h before being washed for 30 sec with 0.3% v/v hydrogen peroxide and thoroughly rinsed with DI water. Fertilized embryos were transferred to large plastic Petri dishes (150mm; Genessee Scientific, Morrisville, NC) containing 25 ppt ASW at a density of ≤50 embryos/dish and kept in a 28 °C incubator with a 14:10 light:dark cycle until exposure.

#### 2.2.2. Killifish: Exposure and Rearing

REPSE was thawed at 4 °C in amber glass jars with PTFE-lined lids just prior to use. Exposure solutions were prepared in glass beakers and contained the specified concentrations (% v/v) of REPSE in 25 ppt ASW. At 24 hours post-fertilization (hpf), killifish embryos were screened for normal development (Armstrong and Child 1965; Oppenheimer 1937). Stage-synchronized embryos from each population were randomly selected and placed individually into 20 mL borosilicate vials (VWR International) each containing 10 mL of exposure solution. Embryos were maintained in a 28 °C incubator with a 14:10 light:dark cycle until use. For individuals undergoing post-hatch endpoints, embryonic REPSE exposure was maintained until 12 days post fertilization (dpf), at which time embryos were removed from exposure solutions and transferred to moist filter paper in a Petri dish. At 14 dpf, hatching was initiated by adding approximately 30 mL of 25 ppt ASW and swirling dishes on an orbital shaker. To synchronize hatching, embryos that had not hatched after 8 h of shaking were treated with 10 mg/mL pronase (Promega, Madison, WI, USA) in ASW for 15 min to soften the chorion sufficiently for the larvae to emerge. Following hatch, larvae were maintained in glass bowls containing approximately 200 mL of 25 ppt ASW and received daily water changes until neurobehavioral assessment at 28 dpf.

Range-finding was conducted using KC embryos, as they are sensitive to PAHs, using concentrations that were selected based on previous studies of ERSE toxicity in KC (Clark et al. 2013; Brown et al. 2016). Controls (0% REPSE) contained only ASW, and the following range of REPSE concentrations were used: 0.05, 0.1, 0.5, 1, 2, 3, 5, and 10%. Range-finding was conducted using the EROD and cardiac teratogenicity assays as described below.

#### 2.2.3. Killifish: EROD Assay

CYP1 activity, a sensitive biomarker of AhR pathway activation by PAHs, was measured using the *in ovo* ethoxyresorufin-O-deethylase (EROD) assay modified from Nacci et al. (1998), as described by Meyer and Di Giulio (2002). Embryos were obtained and exposed for range finding as described above (section 2.2.2), with 5-10 embryos exposed per REPSE concentration across 2 or 3 breeding events for a total of n=15-30 embryos per group. For subsequent comparisons between populations, 5-10 embryos per population per treatment were exposed across 3 breeding events for a total of n=25 embryos per group. In addition to REPSE, exposure solutions for embryos undergoing the EROD assay also contained 21 μg/L ethoxyresorufin in DMSO, where the DMSO concentration in exposure solutions was <0.003%. To assess EROD activity, 96 hpf embryos were imaged using a fluorescence microscope (BZ-X710, Keyence Corporation, Osaka, Japan). For each embryo, fluorescence intensity in the urinary bladder and yolk sac were measured using ImageJ 1.52a (Schneider et al. 2012), with yolk serving as the background correction fluorescence value. EROD activity was calculated for each individual by dividing its background-corrected fluorescence value by the mean of KC control embryos’ fluorescence and then multiplying by 100.

#### 2.2.4. Killifish: Cardiac Teratogenicity Assessment

Embryos were obtained and exposed for range finding as described above, with 5-10 embryos exposed per group across 2 or 3 breeding events for a total of n=15-30 embryos per group. For subsequent comparisons between populations, 5-10 embryos per population per treatment were exposed across 3 breeding events for a total of n=25 embryos per group. At 144 hpf, embryos were observed for cardiac development according to Matson et al. (2008) and Clark et al. (2010). Embryos were scored blind under light microscopy (SMZ1500, Nikon Instruments Inc., Melville, NY, USA) for cardiac teratogenesis. Each embryo was scored as either 0 (normal), 1 (moderate deformity), or 2 (severe deformity). Moderately deformed hearts had a range of alterations from including atrio-ventricular misalignment, elongation of the heart, and/or some pericardial edema. Severely deformed hearts exhibited severe misalignment of the chambers, tube heart, and large pericardial edema.

#### 2.2.5. Killifish: Neurobehavior

To assess neurobehavior, larval locomotor activity was assessed at 18 dpf (6 days post-exposure) and at 28 dpf (16 days post-exposure). Larvae were placed into individual wells of either a 48-well plate (for 18 dpf larvae) or a 24-well plate (for 28 dpf larvae) containing 25 ppt ASW. Following transfer, larvae were acclimated to the plate for 1 h in a dark incubator at 28 °C. Following acclimation, the plate was transferred to the DanioVision (Noldus Information Technology, Leesburg, VA, USA) instrument and maintained at a 28 °C, where EthoVision XT14 tracking software (Noldus Information Technology) was used to track larval movements. Larvae were first habituated in the dark for 10 min followed by 40 minutes of alternating 10 min:10 min light:dark phases. Resulting data were filtered to remove individuals with < 96% quality scores (subject not found in ≥ 4% of frames). For each age, total distance traveled (mm) was assessed for 5-8 larvae (18 dpf) or 2-4 larvae (28 dpf) per group across 3 breeding events, for a total sample size of 12-22 (18 dpf) or 6-13 (28 dpf).

#### 2.2.7. Killifish: Statistical Analysis

Statistical analysis and data visualization for range-finding experiments, cardiac deformity assessment, and EROD were performed in GraphPad Prism 11.0.2 (GraphPad Software, Inc., La Jolla, CA). For range finding, a Shapiro-Wilk was used to test for normal distribution, and significance was tested using Kruskal-Wallis with Dunn’s post-hoc test. For validation of the selected dose, a two-way ANOVA with Dunn-Šidák post-hoc test was used. Statistical analyses for locomotion were performed in JMP 18 (SAS Institute, Cary, NC), and data were visualized in R (R Core Team 2021). Locomotor activity was analyzed using a two-way ANOVA to determine significant differences in total distance traveled during individual light and dark phases (Light Total, Dark Total) and across phases. Dunnett’s post hoc test was used to determine significant difference from KC 0% REPSE (Control). For 28 dpf locomotion, Student’s t test was used to evaluate significant differences in total distance traveled between REPSE treatments within phase for each population.

### 2.3. Zebrafish (*Danio rerio*) Methods

#### 2.3.1. Zebrafish: Husbandry and Breeding

Adult EkkWill wildtype (EkkWill Waterlife Resources, Ruskin, FL, USA) and transgenic *Tg*(cyp1a:NLS-eGFP) (Kim et al. 2013) zebrafish were maintained in a recirculating AHAB system (Pentair Aquatic Ecosystems, Apopka, FL) at a temperature of 28 °C, pH 7.2-7.4, and a 14:10 light:dark cycle. Fish were fed twice daily with freshly hatched *Artemia* nauplii (90% GSL strain; Reed Mariculture, Campbell, CA) in the mornings and Zeigler’s Adult Zebrafish Complete Diet (Pentair Aquatic Ecosystems) in the afternoons.

Static breeding tanks (0.8 L; Pentair Aquatic Ecosystems) were set up in the late afternoon, each with 3 females and 2 males randomly selected from the colony and spawned the following morning. Embryos were collected and pooled, rinsed with DI water, transferred to Petri dishes containing 30% Danieau’s medium (17.4 mM NaCl, 0.21 mM KCl, 0.12 mM MgSO_4_, 0.18 mM Ca(NO_3_)_2_, and 1.5 mM HEPES, pH 7.2) and placed in an incubator at 28 °C until exposure. All procedures involving the care and use of zebrafish were approved by and conducted in accordance with Duke IACUC protocol A069-22-04-25.

#### 2.3.2. Zebrafish: Exposure and Rearing

*Tg*(cyp1a:NLS-eGFP) embryos were used for the *cyp1a* induction assay (section 2.3.3), and EkkWill wildtype fish were used for neurobehavior (section 2.3.4). Embryos were visually inspected for viability and normal development (Kimmel et al. 1995) using a dissecting microscope (Nikon SMZ1500, Nikon Instruments). Normal embryos at 6 hpf (shield stage) were randomly selected and transferred to glass Petri dishes (VWR International) for exposure, with each dish containing 1 embryo/mL. Fish were observed daily for survival and hatching. Hatching was defined as the percentage of surviving individuals in the dish that were hatched. At 96 hpf, they were evaluated for developmental deformities. Fish were then returned to the incubator until they were assessed for *cyp1a* induction or locomotor activity as described below.

#### 2.3.3. Zebrafish: cyp1a expression assay

*Tg*(cyp1a:NLS-eGFP) is a transgenic zebrafish *cyp1a* reporter (EGFP) construct containing all dioxin response elements and cis-regulatory elements required for endogenous expression of *cyp1a* (Kim et al. 2013). Embryos were exposed to REPSE at 6 hpf as described above (section 2.3.2). REPSE was diluted with 30% Danieau medium to produce test concentrations of 0 (Control), 0.5, 1, 5, and 10% for assessments of survival, hatching, and development as described above (section 2.3.2). Results from these observations were used to refine REPSE concentrations for the *cyp1a* expression assay to 0 (Control), 0.001, 0.01, 0.05, 0.1, or 1 %. Experiments were performed across three breeding events with n=30/group in each event (total n = 90/group). At 72 hpf, larvae were transferred to the wells of a black, clear bottom 96-well plate (Corning, Durham, NC, USA), with one larva per well. Larvae were imaged with a fluorescent microscope using a 470/40 nm (em) and 525/50 nm (ex) filter set (BZ-X700 automated microscope, Keyence Corporation of America). Two images were captured for each fish, one fluorescent (EGFP) and one brightfield.

Images were analyzed using Fiji ImageJ 1.53u software (Schindelin et al. 2012). For each EGFP image, the outline of the fish was traced and the fluorescence intensity measured. A background measurement was taken by measuring the fluorescence intensity of a randomly selected portion outside of the fish. These background measurements were subtracted from fluorescence intensity measured in each fish.

#### 2.3.4. Zebrafish: Neurobehavior

EkkWill wildtype zebrafish were exposed and reared as described above (section 2.3.2). REPSE was diluted with 30% Danieau medium to produce test concentrations of 0 (Control), 0.5, 1, and 5% for assessments of survival, hatching, and development as described above (section 2.3.2). All fish exposed to 5% REPSE were severely deformed by 96 hpf and were, therefore, not used for tests of locomotor activity. At 144 hpf, larvae were rinsed of their exposure media by transferring them into clean 30% Danieau medium. They were then transferred to the wells of a 96-well plate, with one larva per well. Plates were acclimated for 1 h in the light at 28 °C and then transferred to the DanioVision (Noldus Information Technology) instrument and assessed for locomotor activity in the same manner as larval killifish (see section 2.2.5). Experiments were performed across three breeding events with n=30-32 individuals per group in each event (total n=93-95 per group).

#### 2.3.5. Zebrafish: Statistical Analysis

Statistical analyses were performed in JMP 18 (SAS Institute, Cary, NC), and data were visualized in R. Survival at 96 hpf and *cyp1a* fluorescence were analyzed by one-way ANOVA with Tukey HSD post-hoc test. Locomotor activity was analyzed using a one-way ANOVA to determine significant differences in total distance traveled during individual light and dark phases (Light Total, Dark Total) and across phases. Dunnett’s post hoc test was used to determine significant differences from control (0% REPSE).

### 2.4. Caenorhabditis elegans Methods

#### 2.4.1. *C. elegans* Maintenance

*C. elegans* strains BY200 (*vtIs1*[p*dat-1*::GFP]), PHX2867 (p*dat-1*::MLS::roGFP), and PHX2923 (p*dat-1*::PercevalHR) were maintained at 20 °C on K-agar plates seeded with OP50 *E. coli* (Boyd et al. 2012). BY200 is a GFP reporter strain for visualization of all dopaminergic neurons, PHX2867 is a roGFP reporter strain for quantification of mitochondrial redox state in dopaminergic neurons, and PHX2923 is a Perceval reporter strain for quantification of the ratio of mitochondrial ATP to ADP levels in dopaminergic neurons. Worms were fed OP50 *E. coli* in experiments requiring culture on solid medium because of the limited bacterial lawn obtained with OP50, allowing easier observation of worms (Stiernagle 2006). HB101 *E. coli* was used in experiments requiring liquid culture because it is less prone to forming bacterial clumps, making it easier to eat for the worms (Avery and Shtonda 2003).

For experiments requiring age-synchronization, a non-starved population of day 1-2 adults was collected in a 15 mL tube from K-agar plates by washing with K-medium (0.5 mL of 10 mg/mL cholesterol, 3 mL of 1 M CaCl2, 3 mL of 1 M MgSO4 per 1 L K-medium) (Sigma-Aldrich, St. Louis, MO, USA; VWR, Radnor, PA, USA) (Boyd et al. 2012). Worms were allowed to settle for 2-3 minutes and the supernatant discarded. Worms were then treated with 5 mL of K-medium containing final concentrations of 0.4 N sodium hydroxide and 20% v/v sodium hypochlorite for eight minutes. The bleaching reaction was quenched by raising the volume to 15 mL with K-medium, centrifuged at 2200 RCF for 2 min, and the supernatant discarded. The last step was repeated two more times to recover the embryos for exposure experiments. This bleaching treatment does not cause dopaminergic neuronal damage in L4 worms (Huayta and Meyer 2025).

#### 2.4.2. C. elegans: Developmental Exposure

Embryos generated by the bleaching treatment were counted on a Leica MZ75 stereomicroscope (Leica, Teaneck, NJ, USA) by transferring 10 µL of K-medium containing embryos to a glass slide. K-medium was added or subtracted until reaching a concentration of 10 embryos/µL. Approximately 100 embryos in 10 µL of K-medium were transferred to each well of a 24-well plate (Greiner Bio-One, Monroe, NC, USA) and the volume was increased to a total of 500 µL per well of complete K-medium containing HB101 *E. coli* bacterial food at a final concentration of OD 2.0, 6 ppt Instant Ocean and either 0, 10, 25 or 50 % v/v of ERSE or REPSE in DI water final concentrations. An additional control well contained no Instant Ocean. The 24-well plate was put on a shaker at 20 °C for 52-54 h to allow worms to reach the L4 larval stage. Afterward, liquid and worms from each well were collected in separate 15 mL tubes, the total volume was raised to 10 mL with K-medium, and the worms were allowed to settle. After 2-3 min, the supernatant was discarded, and this washing step was repeated two more times. At this point worms were ready for endpoint measurement.

#### 2.4.3. C. elegans: Parental Exposure

Exposure was performed as previously described (Huayta et al. 2025) with modifications to account for the use of ERSE and REPSE. Embryos generated by bleaching treatment were transferred to complete K-medium and allowed to hatch overnight. L1 larval stage worms were collected the next day and transferred to K-agar plates seeded with OP50 *E. coli* and left for 48 h to reach their L4 larval stage. These worms were transferred to 24-well plates containing each 500 µL of complete K-medium with HB101 *E. coli* at OD 2.0, approximately 100 L4 stage worms, 6 ppt Instant Ocean and either 0, 10, 25 or 50 % v/v of ERSE or REPSE in DI water final concentrations. An additional control well contained no Instant Ocean. After 24 hours, this parent population was collected by transferring the well content to a 15 mL conical tube. The volume was raised to 15 mL with K-medium. After allowing worms to settle for 5 min, the supernatant was vacuumed, and this washing step was repeated two more times. Worms were then treated with bleach as described in the previous section to recover their embryos. The progeny population was transferred to plates with OP50 *E. coli* for 52-54 h to allow them to reach the L4 stage. At this point, worms were ready for endpoint measurement.

#### 2.4.4. C. elegans: Quantification of developmental delay

L4 worms that were exposed to REPSE during development were collected in 15 mL K-medium and allowed to settle for 5 minutes. The supernatant was discarded and worms were then transferred to unseeded K-agar plates, allowing the worms to dry and disperse on the plate for 5 minutes. Plates containing worms were mounted and imaged using a Keyence BZ-X710 microscope using a Nikon 4X objective in brightfield mode. Images were analyzed using the WormSizer add-in in ImageJ following their published protocol (Moore et al. 2013) to determine the length of 50-100 individual worms per concentration and replicate.

#### 2.4.5. C. elegans: Quantification of neuronal damage

L4 larval stage worms that previously went through either developmental or parental chemical exposure were transferred to 5 µL of 100 mM sodium azide solution on a 2% agarose pad on top of a glass slide. Worms were paralyzed by the sodium azide after one minute and covered with a coverslip. The glass slide was mounted on a Keyence BZ-X710 microscope equipped with a Keyence BZ-X700E metal halide light-source. Z-stacks of individual worms’ heads were acquired using a Nikon 40X objective and a Chroma GFP filter cube with 100 ms of exposure, microscope objective’s pitch of 0.5 µm, and 3×3 binning. Maximum projections of the z-stacks based on maximum intensity were generated, and each dendrite of the cephalic neurons was blind-scored as previously described (Bijwadia et al. 2021). Briefly, 0 – no damage, 1 – irregular (curves), 2 – less than 5 blebs, 3 – 5 to 10 blebs, 4 – more than 10 blebs and/or breaks, 5 – breaks, 25 to 75% dendrite loss, and 6 – breaks, more than 75% dendrite loss.

#### 2.4.6. C. elegans: Quantification of glutathione redox tone and ATP:ADP ratio in dopaminergic neurons

Worms of strains PHX2867 and PHX2923, expressing mitochondrial-localized roGFP and PercevalHR respectively under the *dat-1* promoter (Morton et al. 2023), were used to quantify the ATP:ADP ratio and glutathione redox tone in dopaminergic neurons. Worms that previously went through chemical exposure were transferred to 5 µL of K-medium on an 8% agarose pad on top of a glass slide and covered with a coverslip. No paralytic chemicals were used because many interfere with the endpoints of interest (Morton et al. 2024). The glass slide was mounted on a Keyence BZ-X710 microscope equipped with a Keyence BZ-X700E metal halide light-source. Two images per head of an individual worm, focused on the cell bodies of the cephalic dopaminergic neurons, were acquired using a Nikon 40X objective, 1 and 2 ms of exposure for the GFP and roGFP filter cubes respectively, and 3×3 binning. The first image was acquired using a 470ex/520em filter, and the second image with a 405ex/520em filter. A custom-made MATLAB script was used to perform feature-based segmentation of the cell bodies in each image, background subtraction, and to determine their mean intensity values (Huayta and Meyer 2025). The script then calculated the mean intensity ratios of 405/470 excitation for mitochondrial roGFP representing oxidized to reduced glutathione, and the mean intensity ratios of 470/405 excitation for PercevalHR representing the ATP to ADP ratio in dopaminergic neurons.

#### 2.4.7. C. elegans: Statistical analysis

MATLAB R2024b (MATLAB 24.2; MathWorks, Natick, MA) was used for all statistical testing and graph generation for *C. elegans* data in this manuscript. Statistical tests and sample size are described in their corresponding figure legends.

## 3. Results

### 3.1. Sediment Extract PAH Composition

The complete PAH composition of REPSE is provided in Table S1. Total PAH content (TPAH) of REPSE was measured at 746.7 µg/L. The most abundant PAH was fluoranthene (136.89 µg/L), followed by pyrene (96.08 µg/L) and phenanthrene (85.65 µg/L), with 27 other unique PAH compounds detected. The two polychlorinated hydrocarbons assessed, pentachlorophenol (PCP) and tetrachloro-1,4-benzoquinone (TCBQ), were below the minimum detection limit. As previously reported in Fang et al. (2014), ERSE has a TPAH content of 4967.5 µg/L, with naphthalene (1694.0 µg/L), phenanthrene (671.6 µg/L) and fluoranthene (437.1 µg/L) as the most abundant PAHs along with 23 additional PAHs.

### 3.2. REPSE Toxicity in Atlantic Killifish (*F. heteroclitus*)

#### 3.2.1. Range finding and deformities

Our initial range finding tested REPSE in the sensitive KC population at concentrations of 0.05%, 0.1%, 0.5%, 1%, 2%, 3%, 5%, and 10% REPSE (0.3734 – 74.6705 ug/L TPAHs). Cardiac teratogenesis first became significantly different from control at 2% REPSE (p = 0.0075) and was observed at all higher concentrations (p < 0.0001), with 100% of KC embryos exposed to 10% REPSE exhibiting severe cardiac deformities (score of 2; Fig. 1A).

**Figure 1.**
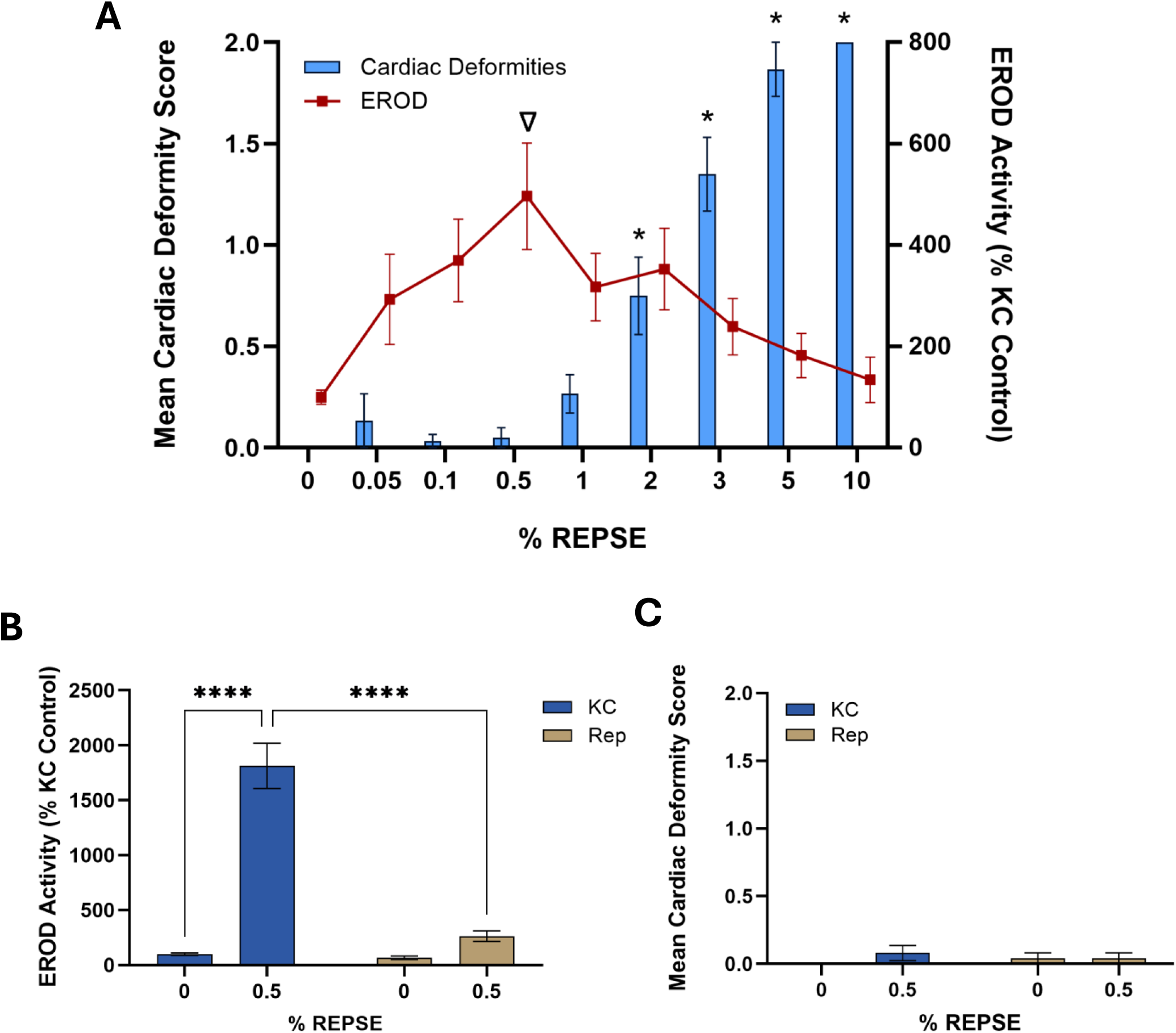
Killifish REPSE range finding and population-specific responses to 0.5% REPSE. **A)** King’s Creek F1 embryo cardiac deformities (blue bars, left axis) and EROD (red line, right axis) after embryonic exposure to REPSE doses ranging 0.05-10%. A Shapiro-Wilk test was used to test normal distribution. Asterisks represent significant difference from control (0% REPSE; *, p < 0.05; Kruskal-Wallis with Dunn’s post-hoc test), and triangle represents chosen test concentration of 0.5% REPSE. Population comparison of KC (blue bars, left) and REP (tan bars, right) responses to embryonic exposure to 0.5% REPSE for **B)** CYP1A induction using the EROD assay and **C)** scoring of cardiac deformities. Asterisks represent significant difference (****, p < 0.0001; two-way ANOVA with Dunn-Šidák post-hoc test).

We also measured CYP1A activity using the EROD assay as a proxy for AhR pathway activation. Only embryos exposed to 0.5% REPSE (3.7335 µg/L TPAHs) showed significantly elevated EROD activity relative to controls (p = 0.0211; Fig. 1A). It should be noted that at higher REPSE concentrations, increasing PAH autofluorescence reduced the sensitivity of the EROD assay after background correction, obscuring additional CYP1A-associated fluorescence despite continued EROD induction (Fig. 1A). However, because these concentrations also produced a high incidence of cardiac deformities, they were excluded from consideration as final test concentrations.

A REPSE concentration of 0.5% was effective at inducing EROD activity in KC without causing significant deformities (Fig. 1A). We therefore repeated the EROD and cardiac deformity assessments in both KC and REP embryos to confirm this concentration as a final test exposure. As expected, PAH-adapted REP embryos showed neither significant CYP1A induction (Fig. 1B) nor cardiac deformities (Fig. 1C) following exposure to 0.5% REPSE. To prevent deformities from confounding neurobehavioral assessments, subsequent experiments initially used only 0% and 0.5% REPSE. A 1% REPSE treatment was added later because, like 0.5%, it induced CYP1A activity without causing significant cardiac deformities (see section 3.2.2).

#### 3.2.2. Neurobehavior

At 18 dpf, population significantly affected total distance traveled (p<0.02, two-way ANOVA), but there was no significant effect of REPSE exposure nor was there a population x REPSE interaction (p>>0.2). REP larvae in the control and 0.5% REPSE groups were significantly hypoactive compared to KC control larvae (p<0.04, Dunnett’s), while REP larvae exposed to 1% REPSE were marginally hypoactive compared to KC controls (p<0.1; Fig. 2A; Fig. S1A).

**Figure 2.**
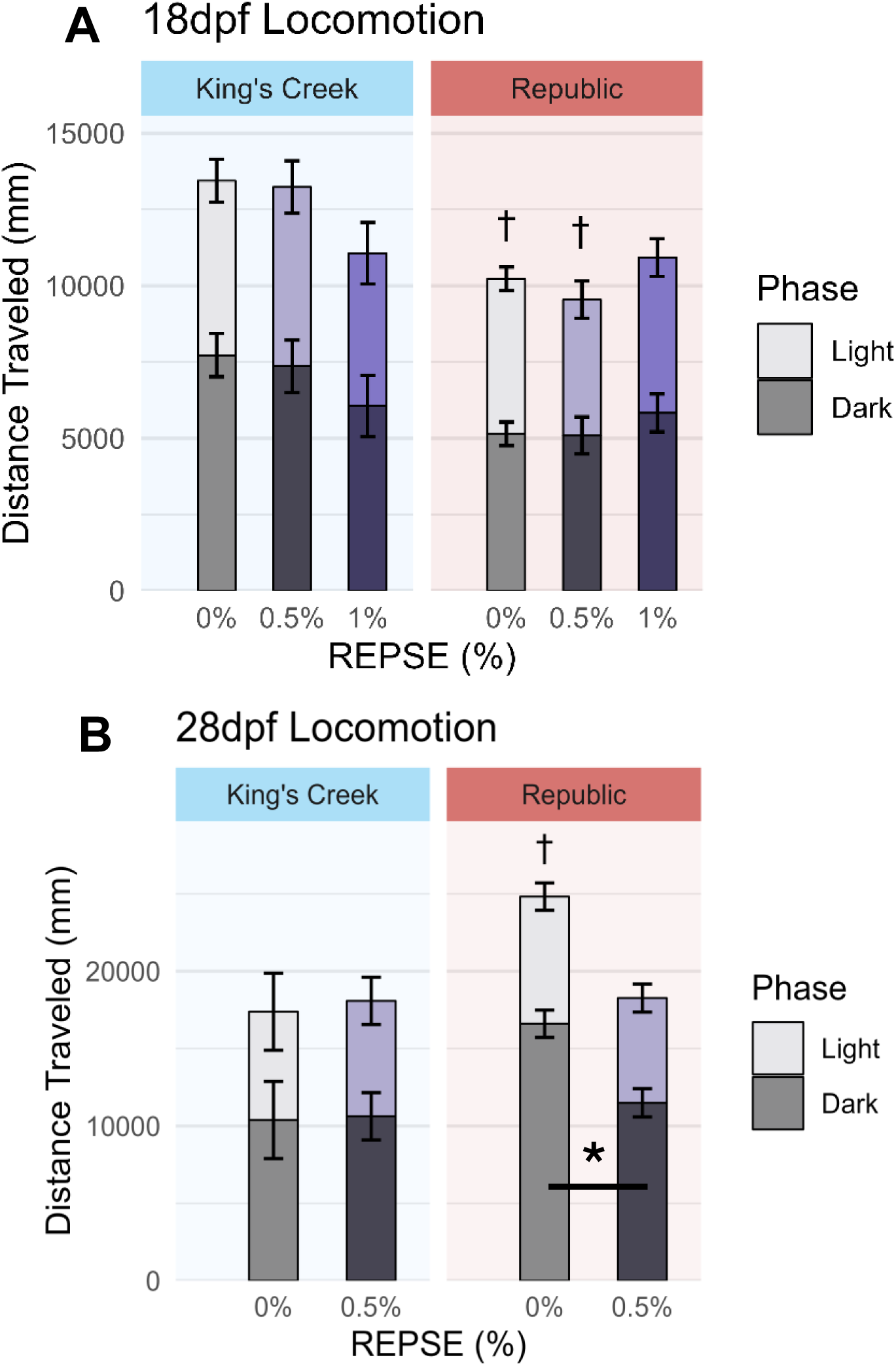
Killifish Neurobehavior after REPSE exposure. Locomotor activity in **A)** 18dpf and **B)** 28dpf killifish larvae. Bars represent means of each phase, where the stacked bar height is the sum of total distance in both phases. Error bars represent SEM within each phase. Daggers represent significant difference of total distance traveled from KC 0% (Control) within age (†, p<0.05; 2-way ANOVA with Dunnett’s post-hoc test). Asterisks represent significant difference between Dark phase distances traveled within population (*, p<0.05; Student’s t).

In contrast, at 28 dpf, REP controls were significantly hyperactive compared to KC controls (p<0.04; Fig. 2B). Interestingly, REP controls were also significantly hyperactive relative to REP larvae exposed to 0.5% REPSE (p<0.03). This difference in REP control larvae was driven by significant hyperactivity in dark phases of the test (Dark phase Population x REPSE interaction, p<0.05; Fig. S1B).

It should be noted that locomotor activity was initially assessed only at 28 dpf using the 0% and 0.5% REPSE treatments. The absence of behavioral effects in KC larvae exposed to 0.5% REPSE, despite significant embryonic CYP1A induction at this concentration (see section 3.2.1), suggested two possible explanations: either 0.5% REPSE was insufficient to disrupt neurodevelopment in KC embryos, or any neurobehavioral effects induced during embryonic exposure had resolved by 28 dpf, sixteen days after exposure ended. To distinguish between these possibilities, we repeated the locomotor assay at 18 dpf and included an additional, higher but still sub-teratogenic concentration (1% REPSE). While we saw a trend towards hypoactivity in KC exposed to 1% REPSE at 18 dpf, suggesting that higher doses of REPSE might impact KC neurobehavior, this was not statistically significant (p>0.3, Dunnett’s).

### 3.3. REPSE Toxicity in Zebrafish (*D. rerio*)

#### 3.3.1. Range finding and deformities

At 96 hpf, we observed a significant decrease in survival of zebrafish exposed to 5% REPSE and 0% survival when exposed to 10% REPSE (Fig. 3A; Fig. S2A). We did not observe significant differences or delays in hatching (Fig. S2B). Developmental deformities that were observed at 96 hpf included craniofacial deformities (*e.g.*, small eyes), pericardial edema, and spinal curvatures (Fig. S2C). All larvae exposed to 5% REPSE exhibited pericardial edema. For this reason, experiments were conducted using sub-teratogenic doses of 1% and lower. REPSE exposure in the *cyp1a* transgenic line, *Tg*(cyp1a:NLS-eGFP), resulted in increased *cyp1a* expression in a dose-dependent manner, with significant differences at 0.05% and higher (Fig. 3B).

**Figure 3.**
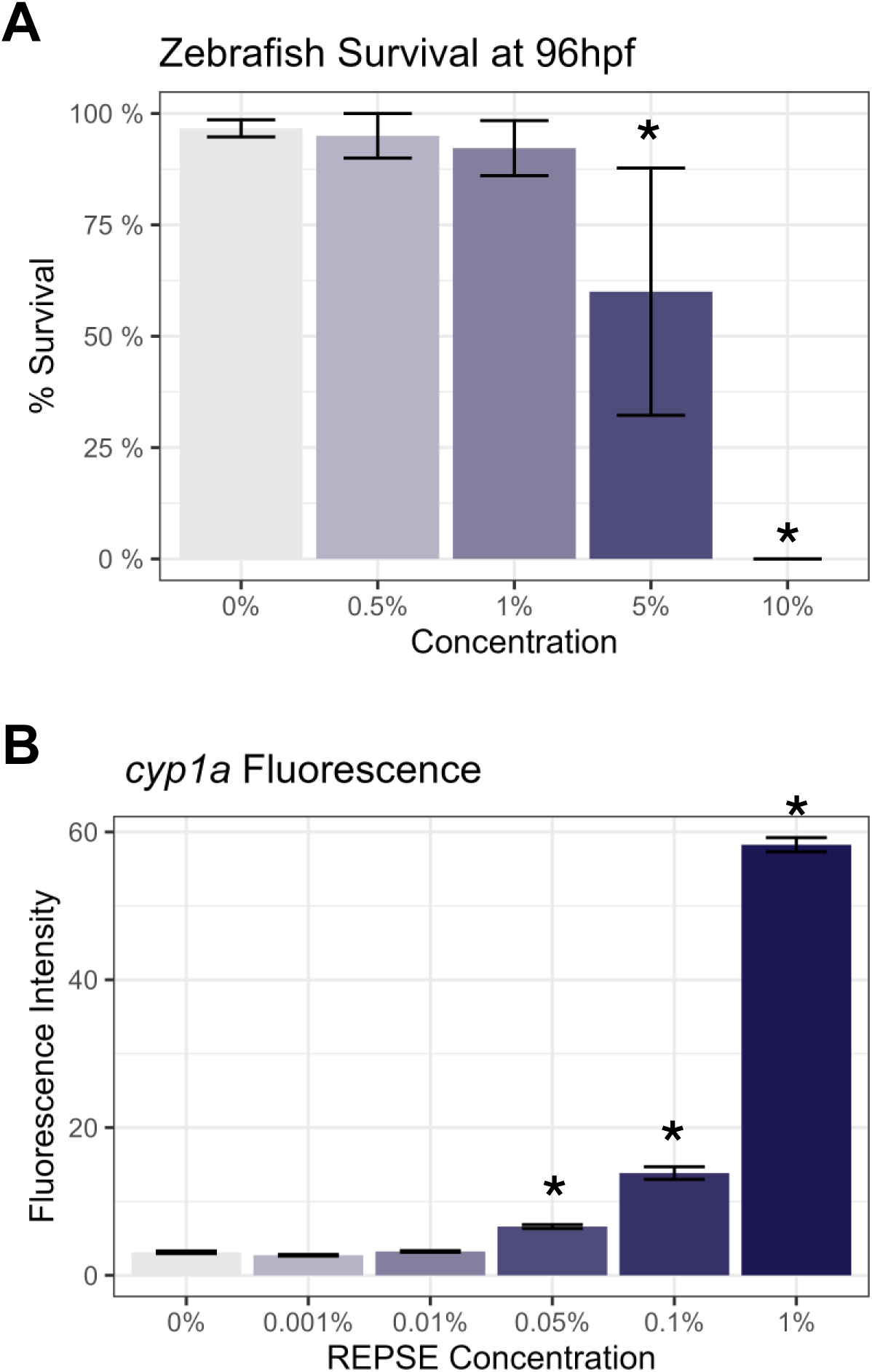
Zebrafish REPSE Range Finding. **A)** 96hpf survival of wild-type EKKWill zebrafish after exposure to a range of REPSE concentrations. **B)** Whole-larvae GFP fluorescence signal (arbitrary units) in 72 hpf *Tg*(cyp1a:NLS-eGFP) zebrafish larvae after exposure to a low range of REPSE concentrations (0.001% - 1%). Columns and error bars represent mean +/− SEM across 3-4 biological replicates. Asterisks represent significant difference from Control (*, p<0.05, one-way ANOVA with Tukey HSD).

#### 3.3.2. Neurobehavior

We observed significant light-phase hypoactivity in zebrafish larvae after developmental exposure to 0.5% and 1% REPSE (Fig. 4; Fig. S3). There were no statistical differences between groups in the dark phases or in total distance traveled.

**Figure 4.**
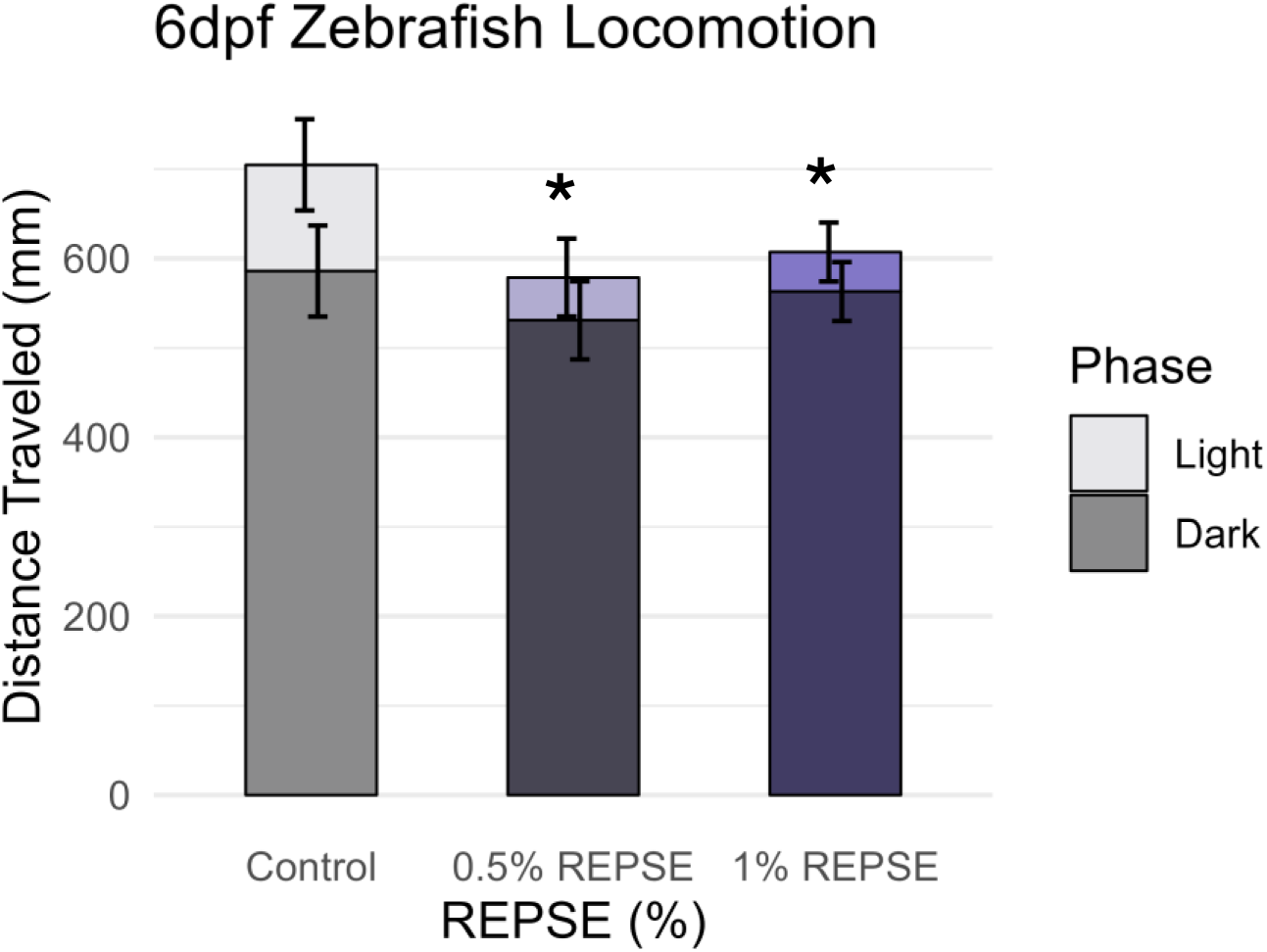
Zebrafish Neurobehavior after REPSE exposure. 6dpf locomotor activity in zebrafish larvae exposed to 0% (Control), 0.5%, or 1% REPSE. **(A)** Sum of total distance traveled by treatment for Light and Dark phases (excluding Habituation from minutes 0-10). Bars represent means of each phase, where the stacked bar height is the sum of total distance in both phases. Error bars represent SEM within each phase. Asterisks represent significant difference between Light phase distances traveled (*, p<0.05; one-way ANOVA with Dunnett’s post-hoc test).

### 3.4. REPSE and ERSE Toxicity in *C. elegans*

#### 3.4.1. Dopaminergic neuronal damage

*C. elegans* were exposed to ERSE and REPSE using both a developmental and a parental exposure paradigm (Fig. 5A). Significant developmental delay after REPSE exposure was observed only for 50% REPSE (p<0.001) (Fig. 5B), indicating the 10% and 25% concentrations were not causative of general organismal toxicity, while 50% led to a reduction of 30% in size with respect to controls. Parental exposure to 10% and 50% ERSE caused significant damage in dopaminergic neurons in offspring, with 10% causing the most pronounced response with respect to the 0% ERSE control (p=0.002) (Fig. 5C). However, no significant effects of parental REPSE exposure were observed in the offspring (Fig. 5D). In contrast, developmental exposures to both REPSE and ERSE caused a significant increase in dopaminergic neuronal damage. This effect was observed at 50% ERSE (p<0.001) (Fig, 5E), and for both 25% (p<0.001) and 50% REPSE (p=0.001) (Fig. 5F).

**Figure 5.**
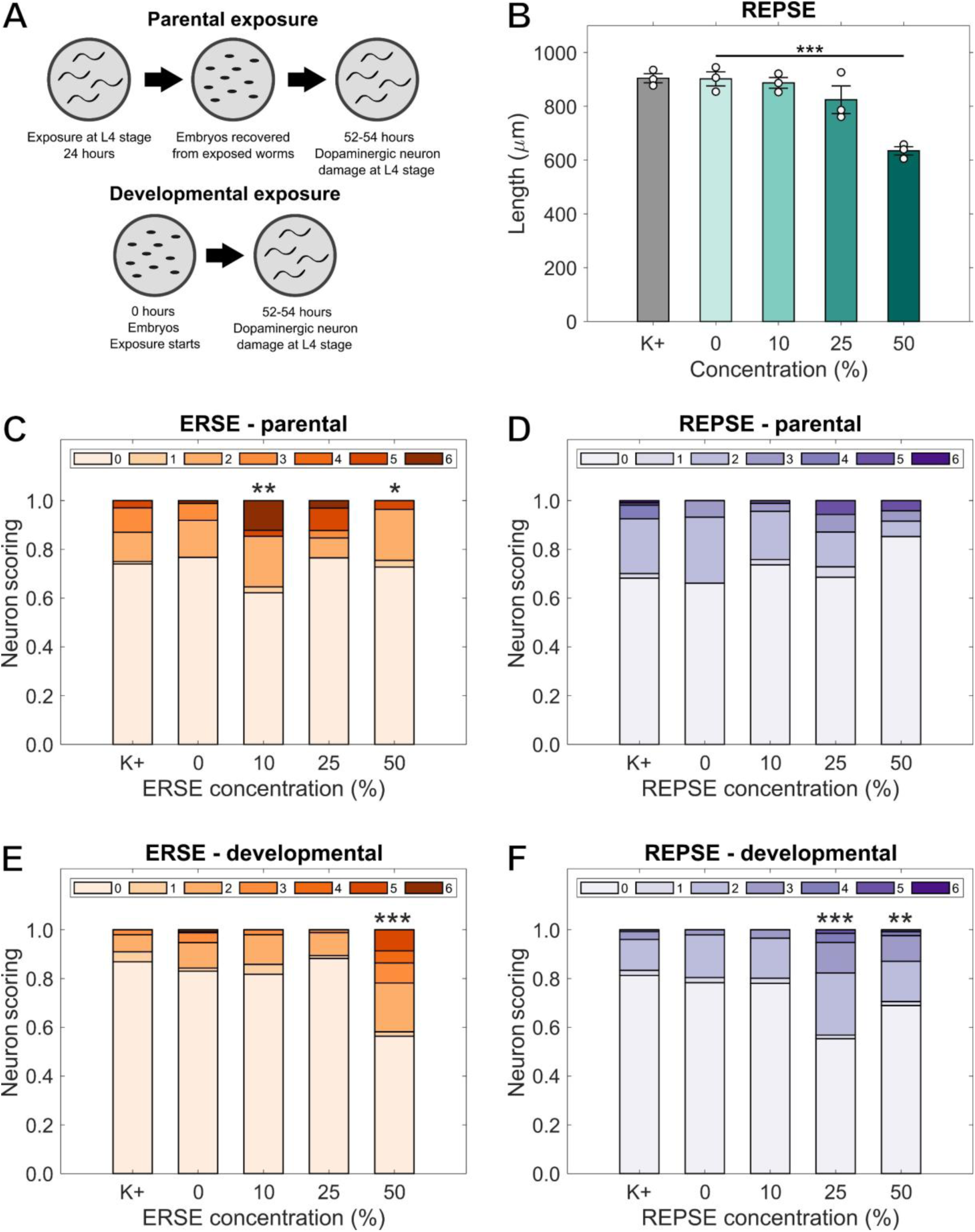
*C. elegans* dopaminergic neurodegeneration after exposure to ERSE/REPSE. **A)** Schematic of parental and developmental exposure paradigms. **B)** Length of worms at the L4 stage after developmental exposure to REPSE N = 3 biological replicates, n = 50-100 worms per replicate per concentration (***, p<0.001; one-way ANOVA with Dunnett’s post-hoc test). Scoring distribution for cephalic dopaminergic neurons of offspring of worms exposed to **A)** ERSE and **B)** REPSE (parental exposure). Scoring distribution for cephalic dopaminergic neurons of worms developmentally exposed to **C)** ERSE and **D)** REPSE (developmental exposure). N = 3 biological replicates, n = 40 dendrites per replicate per concentration. Asterisks represent significant difference (*, p < 0.05; **, p < 0.01; ***, p < 0.001; chi-square with Bonferroni correction).

#### 3.4.2. Redox tone and ATP:ADP ratio

To test whether damage quantified after developmental exposure in dopaminergic neurons was linked to mitochondrial disruption assessed as altered redox state or decreased energetic availability, we repeated developmental exposures with reporter strains for dopaminergic roGFP and ATP:ADP ratio. Both developmental exposures to ERSE and REPSE increased the oxidized to reduced roGFP ratio, a proxy for GSH/GSSG ratio, in dopaminergic neurons. This increase was significant for 10 (p=0.033), 25 (p=0.003) and 50% (p= 0.003) ERSE (Fig. 6A), and for 50% REPSE (p=0.038) (Fig. 6B). However, similar exposures led to a decrease in ATP:ADP ratios only for 50% ERSE (p=0.036) (Fig. 6C) and 50% REPSE (p=0.020) (Fig. 6D) with respect to their 0% controls. Similar quantification after parental exposure did not cause significant changes in roGFP or ATP:ADP ratios for ERSE or REPSE (Fig. S4).

**Figure 6.**
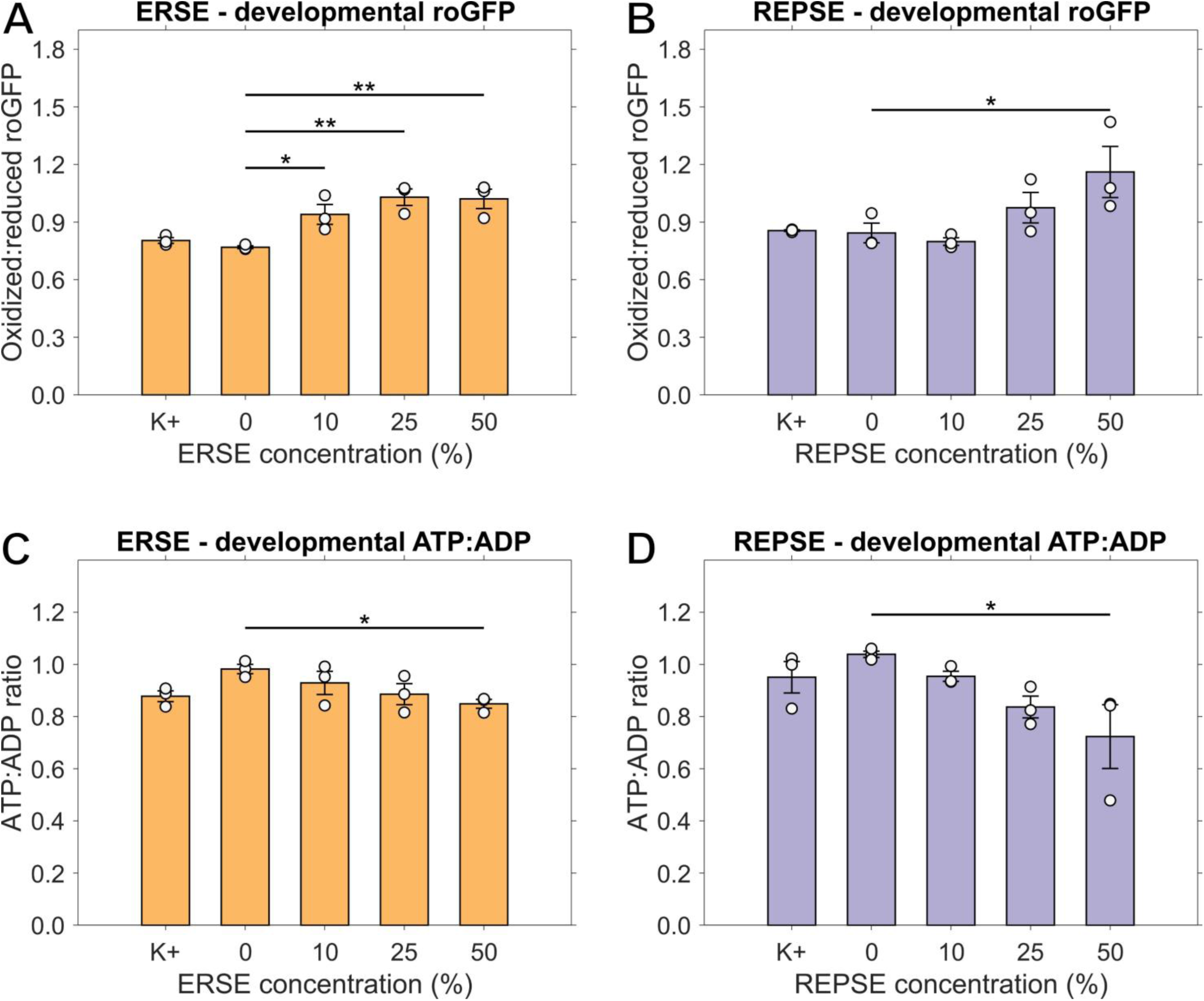
*C. elegans* redox state and ATP levels after developmental exposure to ERSE/REPSE. Oxidized to reduced roGFP ratio in cephalic dopaminergic neurons of worms developmentally exposed to **A)** ERSE and **B)** REPSE. ATP to ADP ratio in cephalic dopaminergic neurons of worms developmentally exposed to **C)** ERSE and **D)** REPSE. N = 3 biological replicates, n = 30-40 neurons per replicate per concentration. Asterisks represent significant difference (*, p < 0.05; **, p < 0.01; 1-way ANOVA with Dunnett’s post hoc test).

## 4. Discussion

### 4.1. Comparison of REPSE and ERSE compositions

The TPAH content of REPSE was approximately 85% lower than that of ERSE (746.7 µg/L versus 5073 µg/L; Fang et al. 2014), a porewater TPAH ratio consistent with previous comparisons of sediment TPAHs at Republic versus Atlantic Wood (Volkoff et al. 2019). The PAH composition of REPSE measured in our study differed from ERSE in the number of individual PAHs found, as well as their relative abundances, thus differing in composition as well as total PAH burden. For example, the top three most abundant PAHs in ERSE are naphthalene (1694.0 µg/L), phenanthrene (671.6 µg/L), and fluoranthene (437.1 µg/L), whereas the top three in REPSE were fluoranthene (136.89 µg/L), pyrene (96.08 µg/L), and phenanthrene (85.65 µg/L). These observations underscore the chemical heterogeneity of environmentally derived PAH mixtures and the limitations of predicting their biological activity based solely on total PAH concentration. Consequently, REPSE and ERSE should not be expected to exhibit directly comparable toxicological potencies or identical mechanisms of action. Rather than directly comparing the relative toxicity of the two mixtures, we used each as a complementary case study to investigate mechanisms of toxicity of environmentally derived PAH mixtures.

### 4.2. REPSE effects on Atlantic killifish

We aimed to gain novel insights into the mechanisms of PAH mixture toxicity by comparing the responses of PAH-sensitive and PAH-tolerant killifish to REPSE exposure. Consistent with previous evidence from complex PAH mixtures, REPSE exposure caused significant teratogenesis in the sensitive KC population. At sub-teratogenic concentrations, despite clear AhR activation as indicated by increased CYP1A activity, KC offspring did not exhibit significant neurobehavioral changes in response to REPSE. Interestingly, the only group in which we observed significant neurobehavioral responses to REPSE was the PAH-adapted REP population. Given the characteristic AhR recalcitrance of REP fish, the neurodevelopmental impacts of exposure to this complex PAH mixture in these killifish are not explained by canonical AhR-mediated mechanisms of developmental PAH toxicity.

Our results provide an unexpected contrast to previous studies using ERSE. Brown et al. (2016) compared behavioral impacts of ERSE exposure between offspring from KC and those from the site of ERSE collection, Atlantic Wood (AW). These authors found a non-monotonic response to ERSE in 18 dpf KC offspring, demonstrating that 0.1% ERSE (5.073 µg/L TPAHs) caused hypoactivity and 1% ERSE (50.73 µg/L TPAHs) caused hyperactivity relative to control. In contrast, the REPSE concentrations we tested in 18 dpf offspring, 0.5% REPSE (3.7335 µg/L TPAHs) and 1% REPSE (7.46705 µg/L TPAHs) did not significantly affect locomotor activity in the same population, KC. Although REPSE contained less TPAHs than ERSE, our results suggest that concentration alone is unlikely to explain these differences.

Another notable difference between the PAH-sensitive and PAH-adapted populations was the locomotor activity profiles of unexposed offspring. In the absence of PAH exposure, the PAH-adapted embryos demonstrated altered neurodevelopment in that REP control larvae were significantly hypoactive at 18 dpf compared to KC control larvae but were hyperactive at 28 dpf. Although previous studies of Elizabeth River killifish did not find significant differences in locomotor activity phenotypes between unexposed offspring from PAH-adapted and PAH-sensitive populations (Brown et al. 2016; Jasperse et al. 2023), pollution-adapted populations of killifish commonly exhibit differences in mitochondrial and bioenergetic phenotypes, which can impact neuronal function (Jasperse et al. 2023; Lindberg 2019).

Our killifish results suggest that Atlantic killifish populations with different pollution exposure histories should not be viewed simply as uniformly “resistant” or “sensitive.” Instead, our findings support a context-dependent framework in which responses to pollutant exposure are interpreted according to contaminant composition and evolved physiological differences among populations. Such an approach may improve our understanding of the mechanisms and evolutionary trade-offs underlying adaptation to chronically contaminated environments.

### 4.3. REPSE effects on zebrafish

The primary goal of using zebrafish in our study was to gain mechanistic insights into REPSE toxicity by comparing neurobehavioral phenotypes with molecular endpoints. Our neurobehavioral assay results demonstrated significant hypoactivity in REPSE-exposed larvae in the light phase, an indication of increased anxiety-like behavior (Hawkey et al. 2022). The use of a *cyp1a* transgenic indicated AhR pathway activation detectable at REPSE concentrations far below those which caused significant neurobehavioral changes.

Although the TPAH concentration in REPSE was approximately one-fifth that of ERSE, the concentrations at which REPSE induced mortality and teratogenesis were not very different from those at which ERSE previously induced toxicity (Fang et al. 2014). This is further evidence that PAH mixture toxicity is not driven simply by the sum total of PAH contents, but by a more complex, composition-dependent mechanism. Past studies comparing individual PAHs have found that their mechanisms of action vary depending on their chemical structure (Incardona et al. 2004, 2006, 2011). Among these, AhR-dependent and cardiac-specific mechanisms have been studied extensively in zebrafish, whereas the molecular phenomena underlying generalized, AhR-independent PAH toxicity remain poorly understood. Early studies put forth the hypothesis that PAHs cause generalized, nonspecific narcotic effects driven by disruption of lipid bilayers (Di Toro and McGrath 2000); however, the non-additive toxicity of PAH mixtures does not fit the additive nature of nonpolar narcosis (Wassenberg and Di Giulio 2004). Given the presence of both AhR-dependent and -independent PAHs in REPSE and ERSE, it is likely that multiple mechanisms are acting concurrently with one another.

### 4.4. ERSE and REPSE effects on *C. elegans*

While both ERSE and REPSE caused significant damage to dopaminergic neurons in *C. elegans*, REPSE caused significant neurotoxicity at different concentrations and did not cause significant damage in offspring after parental exposure. This can likely be attributed to the mixtures having different amounts of total PAHs as well as different distributions of individual components. Notably, *C. elegans* lacks cytochrome P450 1A1 (CYP1A1), leading to limited detoxification pathways for PAHs (Harris et al. 2020). This difference may also partially account for the differences in neuronal damage observed between ERSE and REPSE, and between developmental and parental exposures. Still, these results are consistent with previous descriptions of parental exposure to the representative PAH benzo(a)pyrene leading to neuronal damage in *C. elegans* (Huayta et al. 2025), and increased cell apoptosis being induced by exposure to a complex mixture of PAHs from Deepwater Horizon crude oil (Polli et al. 2020).

Previous evidence connects alterations to morphology of dopaminergic neurons in *C. elegans* with disruption of mitochondrial redox and bioenergetic homeostasis (Morton et al. 2023, 2025). Therefore, we tested for altered mitochondrial redox state or ATP:ADP ratios in dopaminergic neurons after ERSE/REPSE exposure. We quantified a significant increase in oxidized to reduced roGFP for developmental exposures to both ERSE and REPSE. However, the effects were more pronounced with ERSE, starting at a concentration of 10%, while REPSE caused significant damage only at 50%. This correlates well with our neuronal damage results. Similarly, we observed decreased bioenergetics at 50% ERSE and REPSE, with a correlated loss of neuronal integrity. Notably, we did not measure any effects on mitochondrial homeostasis in offspring after parental exposure to ERSE or REPSE, despite quantifying a significant effect in neuronal damage after ERSE parental exposure. REPSE had no effect on neuronal damage after parental exposure, consistent with the lack of change in roGFP or ATP:ADP levels. Possibly, ERSE’s specific composition only affects early neurodevelopment of worms, with acclimation and/or decrease in chemical burden (*e.g.*, due to non-enzymatic excretion or dilution with growth) accounting for the lack of differences in roGFP or ATP levels after parental exposure. Interestingly, we observed a non-monotonic response to parental ERSE for neuronal damage. We speculate that this may be explained by interactions of the chemicals present in ERSE facilitating higher uptake or combinatorial toxicity at lower concentrations. Overall, our results show reduced ATP:ADP ratios and more oxidized roGFP in worms exposed to REPSE during development, but not offspring of exposed worms. Given the lack of PAH-inducible AhR signaling and CYP1A activity in *C. elegans*, the association of REPSE- and ERSE-induced dopaminergic neuronal damage with differential cellular energy availability and increased oxidative stress supports an AhR-independent mechanism of PAH mixture neurotoxicity in this model.

### 4.5. Cross-species insights

By integrating data from model species with complementary strengths, we sought to provide insights into the conserved molecular responses underlying PAH mixture toxicity. PAHs are well known to cause oxidative stress; in vertebrates, this is typically associated with the upregulation of ROS-producing CYP enzymes induced by AhR pathway induction (Knecht et al. 2013; Incardona et al. 2011; Van Tiem and Di Giulio 2011). However, we observed altered redox status in *C. elegans*, which lack CYPs, after REPSE and ERSE exposure. In addition to oxidative stress, we also observed altered cellular energy availability, which is tightly connected to redox status *via* mitochondrial function, in *C. elegans* dopaminergic neurons. Early development is a critical window for the nervous system; disruptions to redox homeostasis and neuronal bioenergetics during this time period are known to cause developmental neurotoxicity (Lin et al. 2020). In this study, developmental REPSE exposure caused later-life alterations to the nervous system in all three models tested, including CYP1A-recalcitrant subpopulations of Atlantic killifish. Taken together, our results support the hypothesis that PAH mixtures can disrupt neuronal bioenergetics through mechanisms that do not require canonical CYP1A-mediated metabolism.

### 4.6. Strengths and limitations of a multi-species approach

Although physiological responses to exposure are often conserved across taxa, they sometimes produce divergent phenotypic outcomes across species and life stages. For example, CYP1A induction remained highly conserved between the two fish species, demonstrating consistent activation of the canonical AhR signaling pathway by REPSE in vertebrates. However, REPSE-induced neurobehavioral phenotypes differed between the killifish and zebrafish, as well as across developmental stages in killifish. These differences likely reflect species-specific behavioral baselines, a product of the ecological contexts that influence how locomotor activity affects evolutionary fitness.

Conversely, while the use of species-specific endpoints complicates direct mechanistic comparisons, our observation of convergent toxicological responses across evolutionarily distant taxa suggests that some key biological pathways underlying PAH mixture toxicity are broadly conserved. REPSE exposure produced neurotoxic effects in all three models, but the endpoints evaluated differed between species. For example, while altered dopaminergic neuronal morphology is correlated with altered behavior in worms (Clark et al. 2024), neuronal damage in *C. elegans* is not mechanistically equivalent to altered neurobehavior in fish; the processes responsible for these phenomena, although they are both governed by the function of the nervous system, are different. Consequently, our mechanistic interpretation across species focuses on broader conserved biological processes rather than strict equivalence of phenotypic outcomes.

## 5. Conclusion

REPSE exposure caused neurotoxicity across three evolutionarily distinct model organisms, and each of the species used in our study provided insights into the mechanisms underlying this. Despite conserved AhR pathway activation by REPSE, as indicated by the induction of CYP1A in both killifish and zebrafish, our killifish results suggested an AhR-independent mechanism of neurotoxicity. In support of this hypothesis, our worm results showed developmental neurotoxicity correlated with altered bioenergetics and redox status, despite the fact that this species’ AhR is not activated by PAHs. Furthermore, by comparing our results with ERSE, a PAH mixture from a nearby site, we found continued evidence that PAH mixture toxicity depends on both concentration and composition. Our findings highlight the importance of mechanistic toxicology approaches that account for both mixture composition and the diversity of biological responses to environmental contaminants. More broadly, these results support the development of mechanistic frameworks that extend beyond the classical paradigms of developmental PAH toxicity.

## 6. Acknowledgements

This work was supported by the National Institute of Environmental Health Sciences (NIEHS) of the National Institutes of Health (NIH) under award numbers P42ES010356 (Duke University Superfund Research Center) and T32ES021432 (Duke University Program in Environmental Health). The content herein is solely the responsibility of the authors and does not officially represent the views of the NIH. We would like to thank Robyn Tanguay’s lab (Oregon State University) for their generous gift of the CYP1A transgenic zebrafish strain, *Tg*(cyp1a:NLS-eGFP), and Michael Aschner’s lab (Albert Einstein College of Medicine) for their gift of the *C. elegans* strain BY200.

