## Supplementary Figures and Table S1 for "Evidence from three taxonomically distinct species for a non-AhR mechanism of developmental neurotoxicity of an environmentally derived mixture of polycyclic aromatic hydrocarbons"

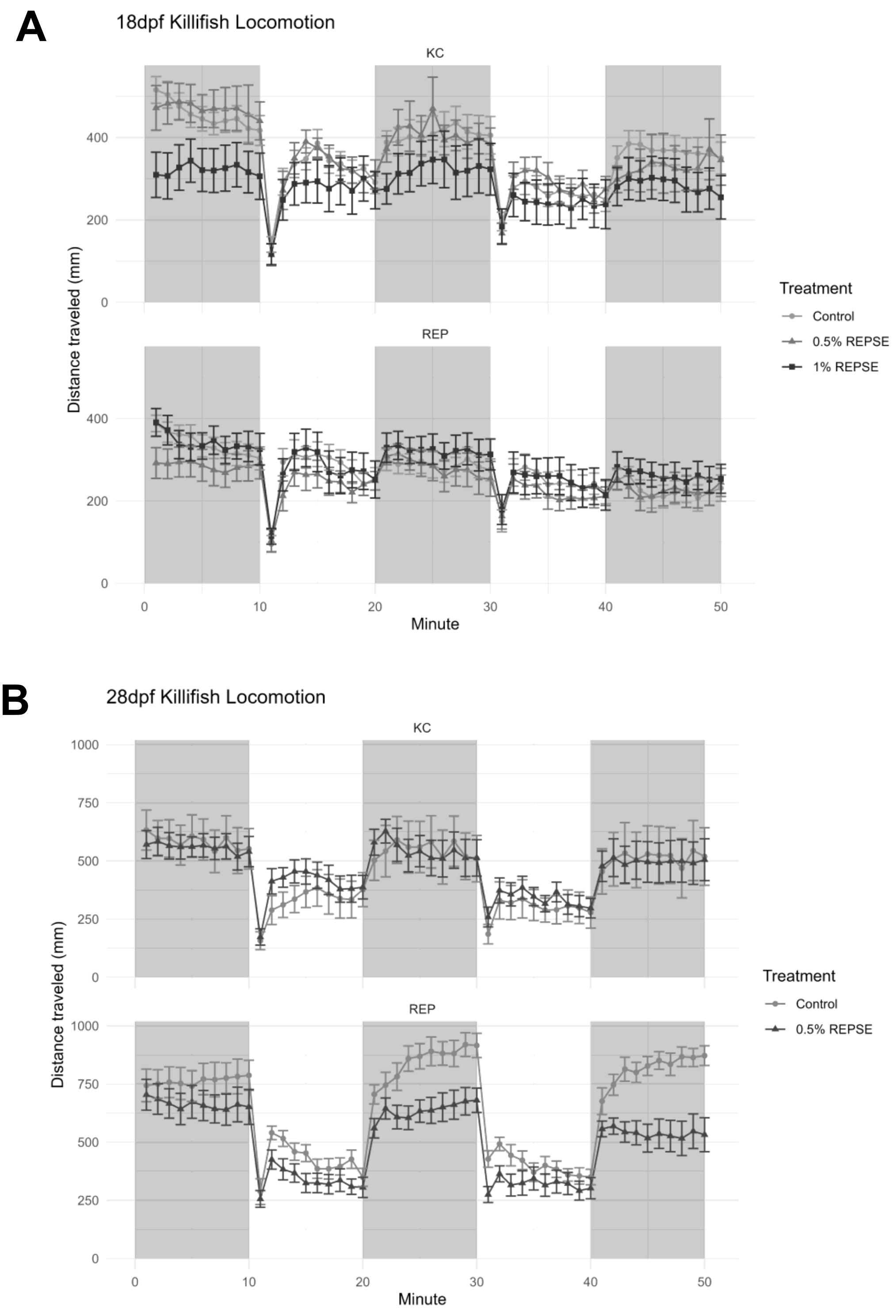


**Figure S1. Killifish Neurobehavior.** Minute-by-minute traces of distance traveled per minute by **A)** 18dpf and **B)** 28dpf killifish larvae from KC (upper graph) and REP (lower graph). Background shading indicates light/dark status of chamber. Points and error bars represent mean +/- SEM for each treatment group.

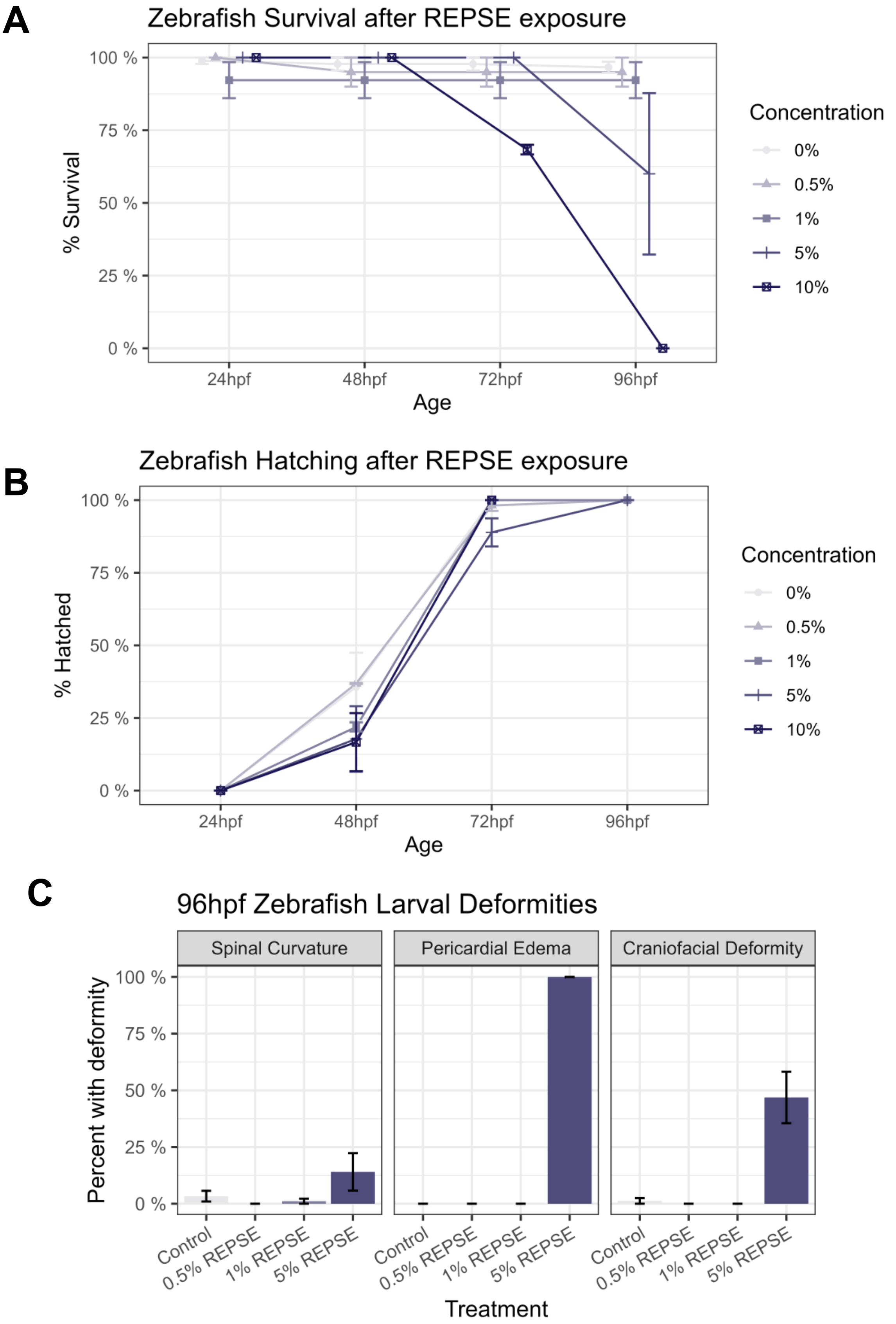


**Figure S2. Zebrafish Survival, hatching, and deformities after REPSE exposure.** Daily survival **(A)** and hatching rates (proportion of surviving individuals hatched) **(B)** for zebrafish larvae from 24-96hpf. Points and error bars represent mean +/- SEM for each treatment group, across 3-4 biological replicates. **C)** Deformity rates by treatment group for frequently observed deformities at 96hpf.

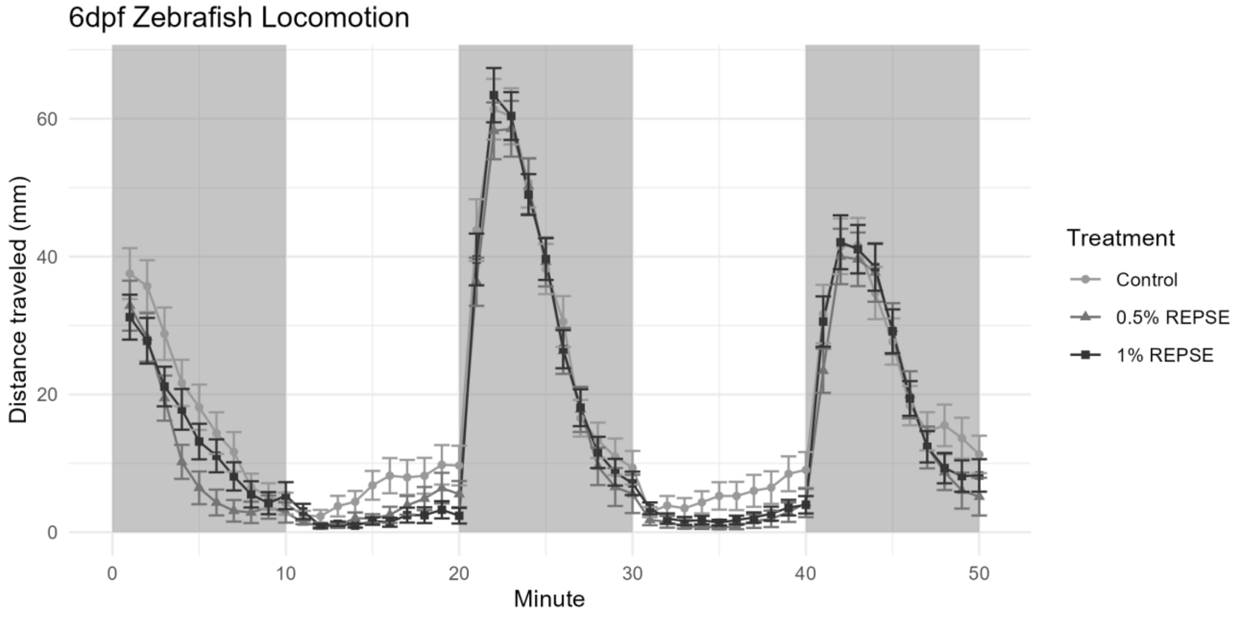


**Figure S3. Zebrafish Neurobehavior.** Minute-by-minute traces of distance traveled per minute. Background shading indicates light/dark status of chamber. Points and error bars represent mean +/- SEM for each treatment group.

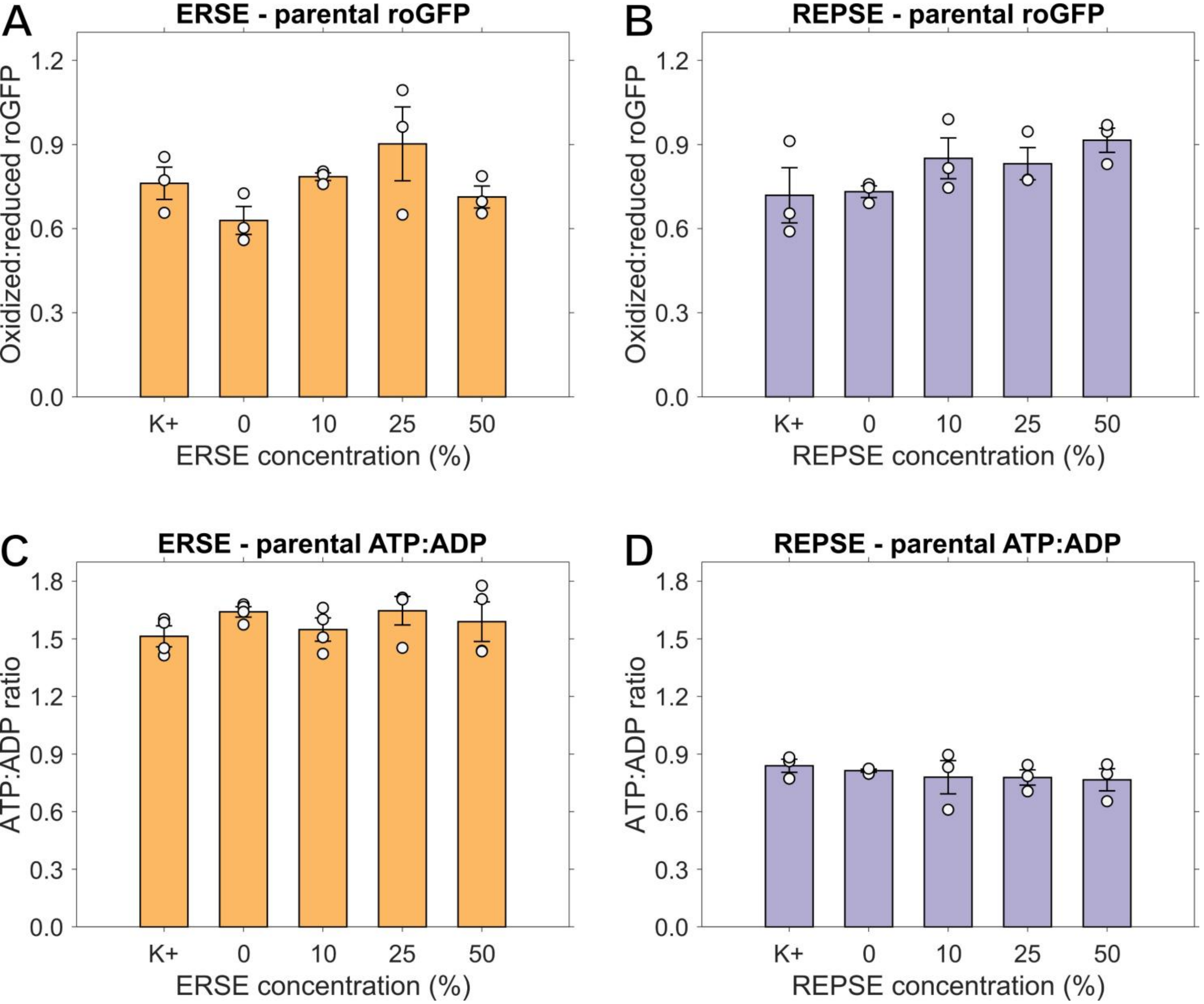


**Figure S4.** ***C. elegans*** **redox state and ATP levels after parental exposure to ERSE/REPSE.** Oxidized to reduced roGFP ratio in cephalic dopaminergic neurons of offspring of worms exposed to **A)** ERSE and **B)** REPSE. ATP to ADP ratio in cephalic dopaminergic neurons of offspring of worms exposed to **C)** ERSE and **D)** REPSE. N = 3-4 biological replicates, n = 30-40 neurons per replicate per concentration, one-way ANOVA with Dunnett's post hoc test.

**TABLE S1.** Concentrations of PAHs from Republic Sediment Extract (REPSE). Values represent averages (ng/mL) of two replicate aliquots of REPSE. Those concentrations below the minimum detection limit are represented by <MDL.

| **Compound** | **Concentration (µg/L)** |
|---|---|
| ***Total PAHs*** | ***746.70*** |
| Fluoranthene | 136.89 |
| Pyrene | 96.08 |
| Phenanthrene | 85.65 |
| Acenaphthene | 47.05 |
| 1,2-Benzanthracene | 40.40 |
| 1,2-benzofluorene | 37.80 |
| Anthracene | 32.83 |
| Fluorene | 26.48 |
| Chrysene | 24.77 |
| Benzo(b)fluoranthene | 22.85 |
| Benzo(a)pyrene | 19.95 |
| Benzo(e)pyrene | 22.01 |
| Benzo(k)fluoranthene | 20.08 |
| Benzo(a)fluoranthene | 16.70 |
| Dibenzofuran | 14.05 |
| 2-methylphenanthrene | 14.29 |
| Carbazole | 11.65 |
| Naphthalene | 10.89 |
| Benzo(b)chrysene | 9.12 |
| 1-methylphenanthrene | 8.69 |
| Indeno(1,2,3-c,d)pyrene | 7.39 |
| Benzo(c)phenanthrene | 8.44 |
| Dibenzothiophene | 6.12 |
| Dibenzo(a,l)pyrene | 5.93 |
| Perylene | 4.52 |
| Dibenzo(a,j)anthracene | 4.57 |
| Acenaphthylene | 2.56 |
| 2,6-dimethylnaphthalene | 2.85 |
| Dibenzo(a,h)anthracene | 2.54 |
| Picene | 1.97 |
| 1-methylnaphthalene | 1.00 |
| Benzo(g,h,i)perylene | 0.52 |
| 3-methylcholanthrene | <MDL |
| Retene | <MDL |
| Pentachlorophenol | <MDL |
| Tetrachloro-1,4-benzoquinone | <MDL |
